# Proteomic analysis of extracellular vesicles released from endothelial cells *in vitro* reveals increased levels of E-selectin and dual specificity phosphatase 7 as a potential marker of TNFα-mediated apoptosis

**DOI:** 10.64898/2026.01.30.702474

**Authors:** Mary E W Collier, Thong Huy Cao, Paulene A Quinn, Jatinderpal K Sandhu, Donald J L Jones, Alison H Goodall

## Abstract

Proteins can be actively packaged into extracellular vesicles (EVs) through mechanisms dependent on the stimulus that activated the cells. Identifying proteins released in endothelial EVs in response to stimuli relevant to cardiovascular disease (CVD) may therefore reveal potential biomarkers that provide information about the vascular endothelium. This study aimed to identify differentially expressed proteins in EVs released from human umbilical vein endothelial cells (HUVEC) in response to stimuli relevant to vascular endothelium activation. HUVEC were stimulated with TNFα (10 ng/mL) or oxLDL (10 µg/mL). Apoptosis was assessed using a flow cytometric DNA fragmentation protocol and caspase-3/7 activity assay. Size distributions of EVs were examined by nanoparticle tracking analysis. Isolated EVs were examined using tandem liquid-chromatography-mass spectrometry (LC-MS/MS). While treatment of HUVECs with TNFα or oxLDL resulted in non-significant elevations in levels of EVs, only TNFα increased apoptosis. Mass spectrometry quantified 1355 proteins and revealed significant differences in the proteome of EVs from TNFα-treated HUVEC compared to EVs from oxLDL-treated or untreated cells. Several candidate biomarkers were significantly and differentially expressed in response to TNFα, including E-selectin and dual specificity phosphatase 7. This study further associated E-selectin on endothelial-derived EVs with endothelial apoptosis and may offer a biomarker of endothelial damage in patients with CVD.

## Introduction

The vascular endothelium plays a key role in maintaining homeostasis by regulating vascular tone [1], releasing anti-inflammatory mediators [2] and providing an anti-thrombotic surface to prevent unnecessary activation of coagulation and inhibit platelet activation [3]. However, processes such as inflammation, changes in shear stress, and oxidative stress can activate and damage vascular endothelial cells, resulting in endothelial activation, endothelial dysfunction and apoptosis [4].

Activation or dysfunction of the vascular endothelium is present in many cardiovascular-related diseases such as atherosclerosis [5], diabetes [6], and renal failure [7]. Interestingly, stimuli that induce endothelial activation and apoptosis such as pro-inflammatory cytokines [8,9,10], low shear stress [11,12], and oxidised low-density lipoprotein (oxLDL) [13] have also been shown to promote the release of extracellular vesicles (EVs) from vascular endothelial cells.

EVs are phospholipid bilayer vesicles that are released from all cell types, including endothelial cells, in response to cellular activation and/or apoptosis. The three main types of EVs include exosomes, which are approximately 30-150 nm in diameter and are derived from the endosomal system, microvesicles (MVs) that bud off directly from the plasma membrane and have a diameter range of approximately 100-1000 nm, and apoptotic bodies, which are larger (1-5 µm) and result from cellular vesiculation during cell death. However, due to significant overlaps in size distributions between different EV populations and the lack of specific markers for exosomes and microvesicles, recent guidelines recommend using the terms small EVs and large EVs unless the source of the EVs can definitely be identified [14]. EVs carry proteins, nucleic acids and phospholipids derived from the parent cell which can be actively packaged into EVs [15,16,17]. Previous studies have revealed that the protein, miRNA and phospholipid composition of EVs is also highly dependent on the stimulus that induces their release, and differs between cellular activation and apoptosis [9,18,19]. This likely reflects the activation of different cellular mechanisms and signalling pathways, resulting in differential loading of proteins and nucleic acids into EVs.

Levels of endothelial cell-derived EVs (ECdEVs) in the blood of healthy individuals is usually low, whereas the presence of increased numbers of ECdEVs in the peripheral circulation has been associated with vascular endothelial dysfunction and apoptosis [20,21,22]. In agreement with this, several clinical studies have detected increased levels of ECdEVs in the peripheral circulation of patients with cardiovascular disease (CVD) compared to healthy individuals [23,24,25,26].

Increased levels of ECdEVs in the circulation of patients with CVD have therefore been suggested as a potential biomarker that could be used for the stratification of patients [26]. Previous studies have used proteomics approaches to examine the protein composition of EVs released from vascular endothelial cells *in vitro* in response to various stimuli such TNFα, high glucose, hypoxia or plasminogen activator inhibitor-1 (PAI-1) [27,28,29,30]. The aim of this project was to use a sensitive proteomics approach to examine in detail the proteome of EVs released from endothelial cells *in vitro* in response to stimuli relevant to dysregulation of the endothelium in CVD, namely TNFα and oxLDL. These stimuli were chosen because of their relevance to CVD and their effects on the vascular endothelium. For example, levels of the proinflammatory cytokine TNFα have been shown to be elevated in the blood of patients with CVD [31,32,33]. TNFα directly affects the vascular endothelium resulting in characteristics of endothelial dysfunction such as increased vascular inflammation and reduced endothelium-dependent vasodilation both in healthy volunteers [34,35] and patients with CVD [36]. Increased levels of circulating oxLDL have also been detected in the circulation of patients with CVD [37,38]. oxLDL induces oxidative stress in vascular endothelial cells resulting in endothelial injury and apoptosis [39]. oxLDL also increases the expression of adhesion molecules on the surface of endothelial cells [40,41] facilitating leukocyte adhesion, and inhibits eNOS activity [42] reducing vasodilation. TNFα and oxLDL were therefore used in this study as stimuli that promote detrimental processes in endothelial cells that may contribute to the development of CVD. As well as examining the proteome of ECdEVs in response to these stimuli, we also compared the protein content of EVs between treatments to establish whether potential biomarkers were specific to different stimuli. This could provide further information about the state of the endothelium in patients with CVD by identifying biomarkers specific to TNFα- or oxLDL-mediated endothelial dysfunction.

## Materials and Methods

### Cell culture

Human umbilical vein endothelial cells (HUVEC) (PromoCell, Heidelberg, Germany) were cultured at 37°C under 5 % (v/v) CO_2_ in Endothelial Cell Growth Medium MV (PromoCell) containing 5 % (v/v) foetal calf serum (FCS). Cells were used between passages 3 and 6.

For experiments, HUVEC were adapted to serum free media (SFM) in three steps. First the cells were incubated at 37°C in MV media containing 2 % (v/v) FCS for 24 h. HUVEC were then washed twice with PBS, pH 7.4 and incubated overnight in SFM containing 0.5 % (w/v) BSA (Sigma/Merck Life Sciences, Gillingham, UK) plus 5 ng/mL recombinant human EGF, 10 ng/mL recombinant human bFGF, 20 ng/mL R3 IGF-1, 0.5 ng/mL recombinant human VEGF 165, 1 µg/mL ascorbic acid, 22.5 µg/mL heparin and 0.2 µg/mL hydrocortisone (PromoCell). Following two further washes with PBS, HUVEC were placed in SFM for 1 h. Finally, the media was replaced with fresh SFM and cells were either left untreated or treated with TNFα (10 ng/mL) (Fisher Scientific, Loughborough, UK) [43,44,45], or a relatively low concentration of oxLDL (10 µg/mL) (Fisher Scientific) [42,46].

### Propidium iodide DNA fragmentation assay

HUVEC (2.5 × 10^5^) were seeded into 6-well plates, adapted to SFM as described above and then treated with different stimuli for 20 h. The media containing floating cells was retained and adherent HUVEC were detached from the plate using TrypLE Select (Fisher Scientific). Floating cells and those detached from the plate were combined and were centrifuged at 800 *g* for 5 min. Cells were then resuspended in HEPES-buffered saline (HBS), pH 7.4 (500 µL) and fixed by the drop-wise addition of 2 mL ice-cold ethanol (70 % (v/v)). Cells were incubated at 4°C for 24 h, followed by centrifugation at 800 *g* for 5 min. Cells were washed with HBS, pH 7.4 (1 mL), resuspended in HBS (200 µL) containing propidium iodide (2 µg/mL) (BD Biosciences, San Jose, US) and DNase-free RNase A (50 U/mL) Fisher Scientific) and incubated at room temperature for 30 min in the dark. The samples were then centrifuged at 800 *g* for 5 min, resuspended in HBS, pH 7.4 (400 µL) and examined on a Gallios flow cytometer (Beckman Coulter, High Wycombe, UK). Cells were analysed by forward scatter vs. side scatter plots and single cells were gated to exclude doublets using peak height against fluorescence on the FL-3 channel. A histogram plot was used to distinguish between cells in different stages of the cell cycle corresponding to apoptotic cells (fragmented DNA), cells in G1 of the cell cycle and cells in G2.

### Caspase 3/7 activity assay

HUVEC (5 × 10^3^) were seeded into 96-well plates in 200 µL of MV media and adapted to SFM. Cells in SFM (100 µL) were treated with different stimuli and incubated at 37°C for 6 h. Caspase-Glo 3/7 assay substrate (100 µL) (Promega, Southampton, UK) was added to each well, mixed on an orbital shaker at 500 rpm for 30 s and then incubated at room temperature for 1 h in the dark. Cell lysates were transferred to white-walled 96-well plates and luminescence recorded using a NOVOstar plate reader (BMG LabTech, Aylesbury, UK). Background luminescence, determined as luminescence in wells with cell culture media without cells, was subtracted from the luminescence values of the sample wells.

### Determination of cell numbers using crystal violet staining

HUVEC (5 × 10^4^) were seeded into 96-well plates, adapted to SFM as described above and then treated with different stimuli and incubated for 20 h at 37°C. Cells were washed with PBS, pH 7.4 (200 µL) and fixed with 4 % (w/v) paraformaldehyde (75 µL) for 7 min at room temperature. Cells were washed three times with PBS and stained with crystal violet (50 µL) for 30 min. Samples were then washed three times with distilled water (200 µL) and 1 % (w/v) sodium dodecyl sulphate (SDS) (50 µL) added per well to elute the crystal violet stain. Absorbances of the samples were measured at 595 nm using a KC Junior plate reader (BioTek, Swindon, UK).

Absorbances were converted to cell numbers using a standard curve of cell numbers against absorbance at 595nm.

### Nanoparticle tracking analysis of EVs

HUVEC (2.5 × 10^5^) were seeded into 6-well plates, adapted to SFM and treated with different stimuli for 6 h as described above. Cell culture media was then centrifuged at 1500 *g* for 15 min to remove cell debris. The cleared media was retained and total EC-EVs were immediately analysed by NTA using a NS500 machine and NTA 2.3 software (Malvern Panalytical Ltd, Malvern, UK). Five 60 s videos were taken for each sample with a camera level of 13 at 25°C. Data was analysed using the NTA 2.3 software with a detection threshold of 10.

### EV isolation from conditioned media for mass spectrometry

HUVEC (2.5 × 10^5^) were seeded into T75 flasks in MV media containing 5 % (v/v) FCS and incubated at 37°C for 72 h. Cells were then adapted to SFM containing 0.5 % (w/v) BSA and growth factors for 24 h as described above. On the day of EV isolation, HUVEC were washed twice with 10 mL of PBS and adapted to SFM (10 mL) for 1 h. The media was then replaced with 10 mL of fresh SFM and cells were either left untreated or incubated with different stimuli and incubated at 37°C for 6 h to allow for the release of EVs. The media was then centrifuged at 1500 *g* for 15 min to remove cell debris, followed by centrifugation at 40,000 *g* for 1 h at room temperature on a Sorvall RC 6 Plus high-speed centrifuge (Fisher Scientific) using an SA-512 rotor (Thermo Scientific) to pellet the EVs. The EVs were washed by resuspension in 8 mL of HBS, pH 7.4 to remove contaminants and centrifuged again at 40,000 *g* for 1 h. The EVs were resuspended in 50 mM ammonium bicarbonate, pH 7.6 (200 µL). Three biological replicates of each group (untreated, TNFα-treated and oxLDL-treated) were prepared as independent experiments and stored at -80°C.

### Preparation of EV samples for mass spectrometry

EV samples in ammonium bicarbonate (200 µL) were reduced using 5 mM dithiothreitol at 65°C for 30 min in the presence of 0.5 % ammonium deoxycholate, pH 8. Samples were alkylated using 20 mM iodoacetamide (IAA) in ammonium bicarbonate, pH 7.6 for 30 min at room temperature in the dark. Proteins were digested using acetylated trypsin from bovine pancreas (5 µL at 1mg/mL; Sigma) at 37°C overnight. Detergents were then precipitated by the addition of formic acid (FA) (1 % (v/v) final concentration) and samples were centrifuged at 20,000 *g* for 5 min.

Samples were loaded onto Sep-Pak C18 cartridges (Waters, Milford, MA, USA) and peptides allowed to bind for 5 min. The columns were washed three times with 0.1 % FA and peptides were eluted in 60 % LC/MS grade acetonitrile (ACN) (600 µL) followed by 80 % ACN (600 µL). Solvents were removed by centrifugation in a SpeedVac concentrator for 1.5 h. Samples were then freeze-dried overnight and resuspended in 20 µL of 0.1 % FA.

### Waters NanoAcquity UPLC and Synapt G2S mass spectrometry

Sample separation was performed using an Acquity UPLC Symmetry C18 trapping column (180 μm x 20mm, 5 μm) to remove salt and any other impurities and a HSS T3 analytical column (75 μm x 150 mm, 1.8 μm). Solvent A was compromised of 0.1 % FA in HPLC grade water and solvent B contained 0.1 % FA in ACN. Supplemental Data File 1 shows the gradient in 110 minutes of solvent A and B used in LC ESI-MS/MS analysis. The flow rate of solvents was 0.3 μL/minute. The Acquity UPLC was coupled to a Water Synapt G2S mass spectrometer (Waters Corporation, Manchester, UK) coupled directly to the Nano Acquity UPLC was used in this study. Analysis using the G2S used HDMSE DIA mode using a Waters Zspray NanoLockSpray in ESI positive mode. The lockspray compound was glu-fibrinogen peptide (GFP), 100 fmol/µL in 50:50 MeCN:H2O + 0.1% FA, measured at an m/z of 784.84265. The mass range was set to 50-2000 Daltons. The IMS wave velocity was 700m/s. Ion mobility-dependent collision energy profiles were used as a .LUT files (Supplemental Data File 2) [47]. The cone voltage was set to 30 v, and the capillary voltage was set to 3.0 kV. Data was acquired using MassLynx V4.2. Each sample was analysed in triplicate.

### Analysis of LC-MS/MS data

Proteomics analysis was carried out using Progenesis QI for Proteomics version 4.2 (Nonlinear Dynamics, Waters Corporation, UK). Spectral data were matched to entries in the UniProtKB human FASTA database for peptide and protein identification. Trypsin was designated as the proteolytic enzyme that allowed up to two missed cleavage sites and maximum protein molecular weight threshold of 1000 kDa was applied. Cysteine residues were modified by carbamidomethylation as a fixed modification while deamidation N, oxidation M, and phosphorylation STY were treated as variable modifications. Identification was considered valid with at least two fragment ions per peptide, five fragment ions per protein, and a minimum of two peptides per protein. The abundance means were calculated at the protein level. The false discovery rate (FDR) was rigorously maintained below 1% for both peptide and protein identifications. Protein quantification was based on the Hi3 approach that utilised the three most intense peptides per protein. The finalised data outputs were then transferred to Microsoft Excel for downstream statistical evaluations and bioinformatics interpretation. Missing values were imputed by replacing the value of minimum/2 for each protein in the matrix from the normalised abundance data. The means of the technical triplicates for each sample were calculated, resulting in three means for each biological replicate for each treatment group. The means were then used to generate fold changes and *p*-values for comparisons between the groups using the Limma R package [48]. *P*-values were calculated between two groups and adjusted for multiple testing errors using Benjamini-Hochberg (BH) with a false discovery rate (FDR) of 0.05. Log2 fold changes (log2FC) were plotted against adjusted -log10 p-values for each comparison with *p*-values <0.05 and log2FC >1 or <-1 considered as significant. STRING protein-protein interaction analysis [49] was used to examine networks of significant DEPs with *p*-values <0.05 and fold changes >2. The highest confidence level (0.9) and evidence for interactions from all sources was used for STRING analysis. Gene ontology (GO)-term enrichment analysis of significant DEPs was examined using the PANTHER overrepresentation test (GO Ontology database DOI: GO Ontology database DOI: 10.5281/zenodo.10536401 Released 2024-01-17) with Fisher’s exact test and false discovery rate (FDR) correction. Only biological process terms and cellular component terms that had an FDR of p<0.05 were included.

## Results

### Treatment of HUVEC with TNFα but not oxLDL increased apoptosis

Flow cytometric analysis of DNA fragmentation was used to assess levels of apoptosis in untreated, TNFα-treated, or oxLDL-treated HUVECs. Propidium iodide-stained cells were examined by forward scatter vs. side scatter plots and single cells were gated to exclude doublets using peak height against fluorescence on the FL-3 channel (Figure 1A). Histogram plots for untreated (Figure 1B), TNFα-treated (Figure 1C) and oxLDL-treated cells (Figure 1D) showed three peaks corresponding to cells in G1 and G2 phases of the cell cycle, and apoptotic cells containing fragmented DNA. Treatment of HUVEC with TNFα (10 ng/mL) for 20 h increased levels of DNA fragmentation and significantly reduced the number of cells in the G1 phase of the cell cycle compared to untreated cells (*p* = 0.0025 TNFα vs. untreated cells), or cells treated with oxLDL (*p* = 0.0007 TNFα vs. oxLDL) (Figure 1E). A significant increase in caspase-3/7 activity was observed following 6 h incubation of HUVEC with TNFα compared to untreated cells (*p* = 0.0018 TNFα vs. untreated cells), or cells treated with oxLDL (*p* = 0.0002 TNFα vs. oxLDL) (Figure 1F). Finally, crystal violet staining revealed a significant reduction in cell numbers in HUVEC incubated with TNFα for 20 h compared to oxLDL (*p* = 0.0342 TNFα vs. oxLDL) (Figure 1G). Treatment of HUVEC with oxLDL (10 µg/mL) had no significant effect on DNA fragmentation (Figure 1E), caspase 3/7 activity (Figure 1F) or cell numbers (Figure 1G) compared to the untreated controls. Taken together these data show that treatment of HUVEC with TNFα resulted in a halt in cell cycle progression and increased apoptosis, whereas oxLDL-treated cells were similar to untreated cells and were not apoptotic.

**Figure 1.**
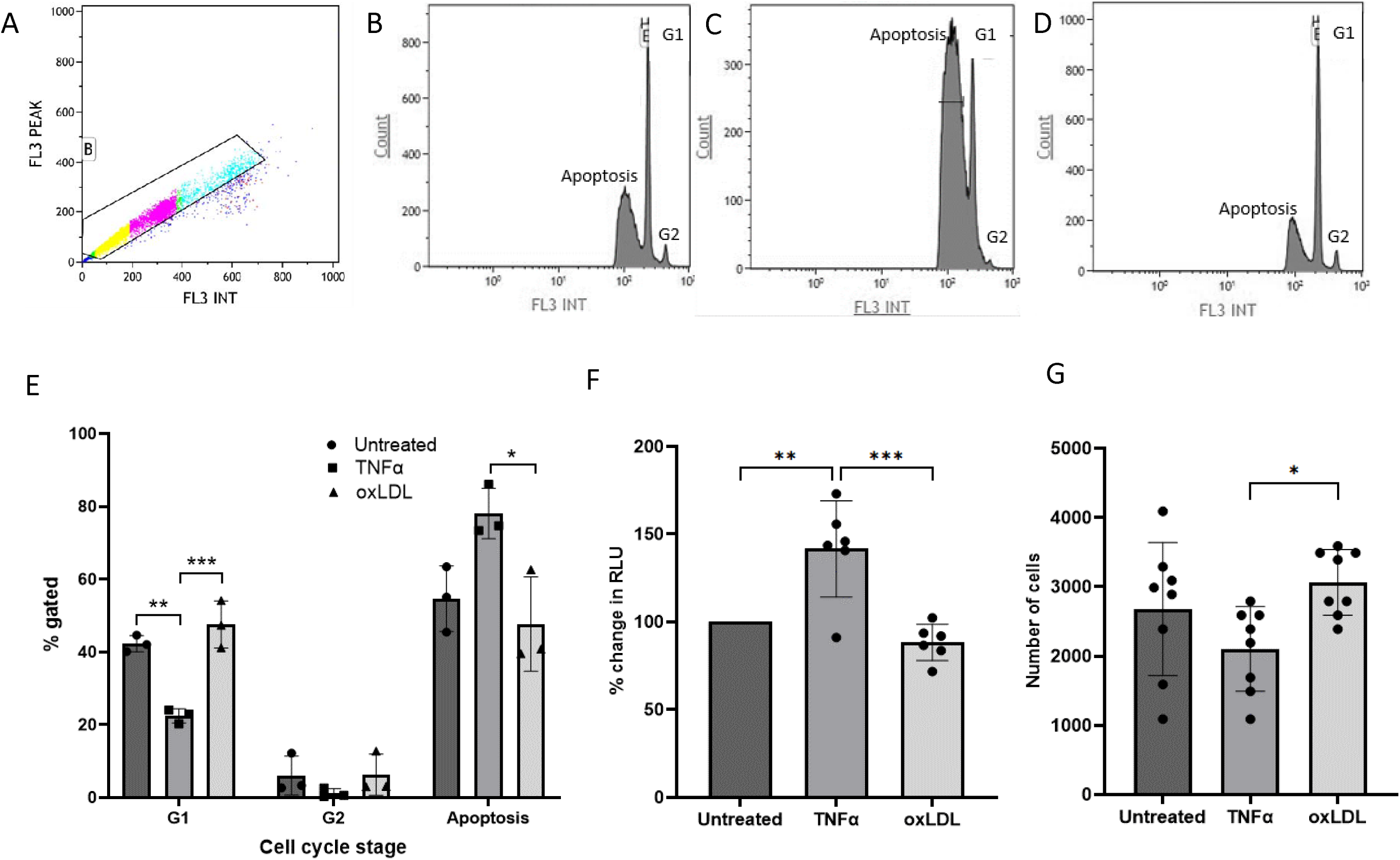
Apoptosis in HUVEC treated with different stimuli. HUVEC were adapted to SFM and treated with TNFα (10 ng/ml) or oxLDL (10 µg/ml) for 20 h. DNA fragmentation was examined by staining cells with propidium iodide staining followed by flow cytometric analysis. (A) Cells were examined by forward scatter vs. side scatter plots and single cells were gated to exclude doublets using peak height against fluorescence on the FL-3 channel. (B) Histogram for untreated cells, (C) TNFα-treated cells and (D) oxLDL-treated cells. Gates were set to define three populations of cells: cells in G1 of the cell cycle, cells in G2, and apoptotic cells with fragmented DNA. (E) The percentage of cells in each gate was calculated (n=3, One Way ANOVA, Tukey’s test ± SD) ** *p* = 0.0025; *** *p* = 0.0007; * *p* = 0.0226. (F) Caspase 3/7 activity was measured following 6 h incubation of HUVEC with different stimuli. Percentage change in RLU compared to the untreated control was calculated for each set (n=6, One Way ANOVA, Tukey’s test ± SD) ** *p* = 0.0018; *** *p* = 0.0002. (G) Crystal violet staining was used to determine the number of cells following treatment with stimuli for 20 h (n=8, One Way ANOVA, Tukey’s test ± SD) * *p* = 0.0342.

### Characterisation of HUVEC-derived EVs

NTA was used to determine the size distributions of ECdEVs in HUVEC conditioned medium following stimulation of cells for 6 h. NTA analysis showed that HUVEC treated with TNFα and oxLDL released similar numbers of EVs, but the oxLDL ECdEVs were generally smaller, with 40.3 % of the released vesicle population in the 30-150 nm size range, compared to 31.4 % of ECdEVs from TNFα-treated HUVEC in this size range. The mean diameter of ECdEVs released from untreated cells was 426 ± 139 nm (Figure 2A), compared to 321 ± 29 nm for the TNFα-treated cells (Figure 2B), and 291 ± 59 nm for the ox-LDL-treated cells (Figure 2C). The total numbers of ECdEVs release in response to TNFα and oxLDL were higher compared to untreated cells, although these increases did not reach statistical significance (Figure 2D).

**Figure 2.**
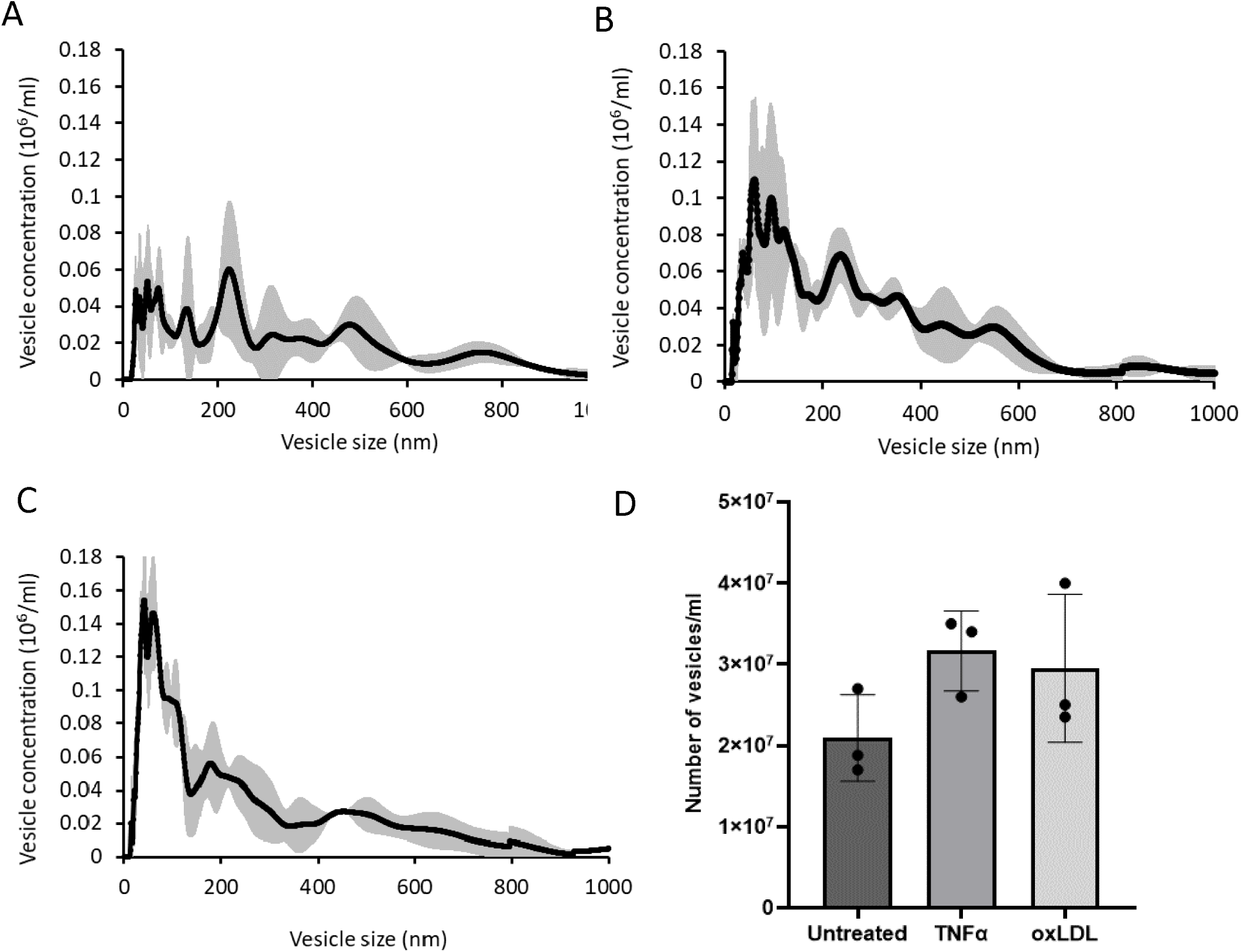
Characterisation of HUVEC-derived EVs. HUVEC (2.5 × 10^5^) in 6-well plates were adapted to SFM and treated with TNFα (10 ng/mL) or oxLDL (10 µg/mL) and incubated at 37°C for 6 h. The conditioned media was then removed and centrifuged at 1500 *g* for 15 min to remove cell debris. The size distributions and concentration of EVs in the media were examined by NTA. Figures show the size distributions of EVs from (A) untreated HUVEC, (B) HUVEC treated with TNFα, (C) HUVEC treated with oxLDL (n=3 ± SD). (D) The concentrations of EVs released in response to different stimuli were also determined using NTA (n=3 ± SD).

### Proteomic analysis of HUVEC-derived EVs confirmed the presence of EV markers

In total, 1784 proteins were detected in EVs released from HUVEC and 1355 were quantifiable. Of the top 100 EV-associated proteins listed in the ExoCarta and Vesiclepedia databases, 90 and 85 (respectively) were present in the HUVEC-derived EVs. These included tetraspanins (CD81 and CD9), flotillin 1 and 2 (FLOT1, FLOT2), integrins (ITGB1, ITGA6), annexins (ANXA1, 5, 6), members of the heat shock protein family (HSPA1A, HSP90AB1, HSPA5), small GTPases (RAB5C, RAB14, RAP1B), and Alix (PDCD6IP) which is involved in multivesicular body formation. This demonstrates a comprehensive coverage of the expected proteins found in EVs.

### Proteomic analysis of HUVEC-derived EVs revealed significantly and differentially expressed proteins dependent on conditions

Since treatment of HUVEC with TNFα or oxLDL resulted in increased EV release compared to untreated cells, we used mass spectrometry to compare the protein content of EVs released from HUVEC in response to these stimuli and EVs from untreated cells. Principal component analysis (PCA) of the mass spectrometry abundance data showed clear discrimination between proteins detected in EVs from TNFα-treated, oxLDL-treated and untreated cells (Figure 3A). Log2FC and adjusted *p*-values for of all quantifiable proteins in EVs for all treatments are shown in Supplemental Data File 3. Additionally, peptide counts, number of unique peptides and confidence scores for all 1784 proteins are shown in Supplemental Data File 4.

**Figure 3.**
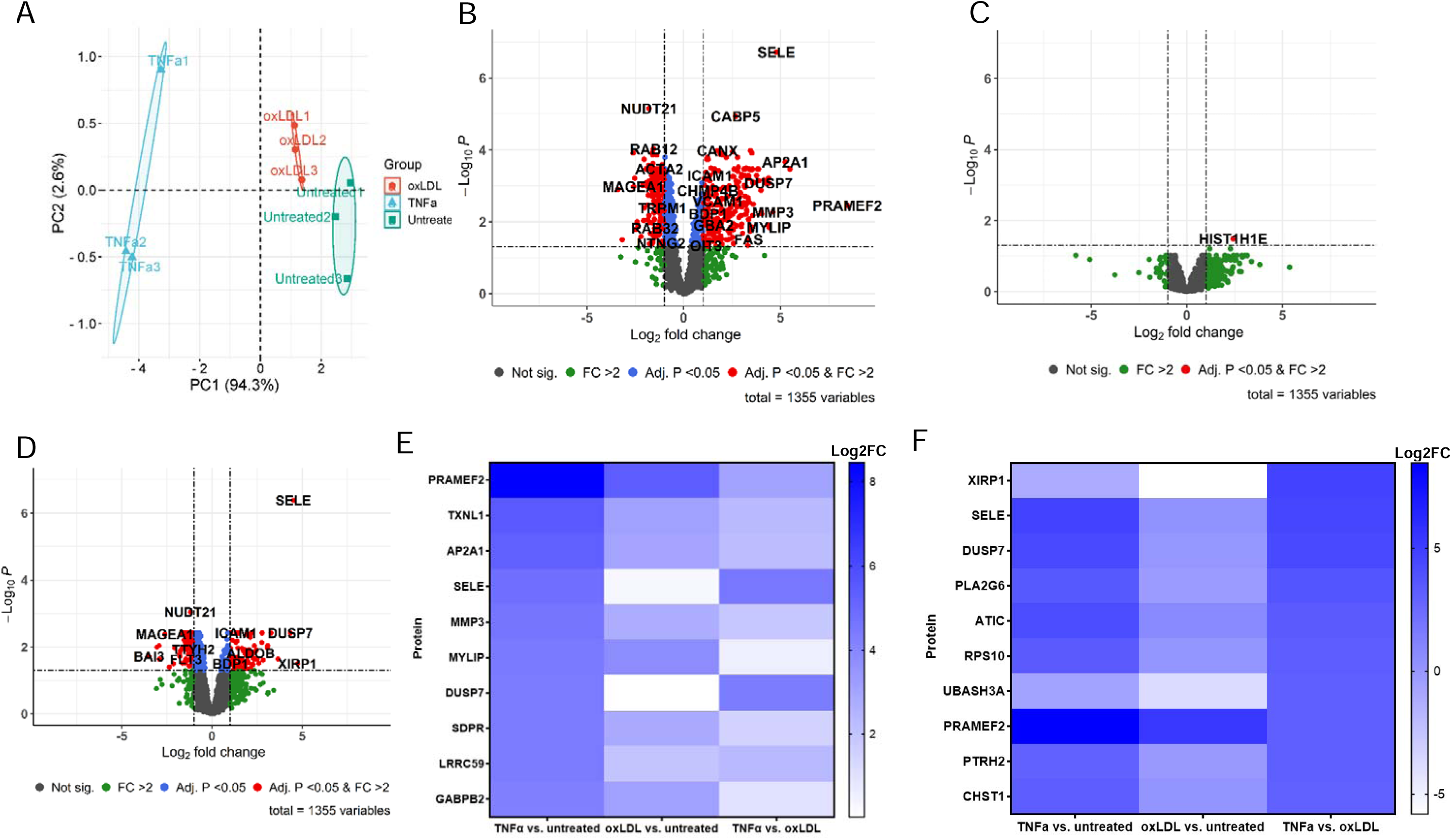
Analysis of proteins detected in EVs in response to treatment of HUVEC with TNFα or oxLDL. A) Principal component analysis showed good discrimination between proteins in HUVEC-derived EVs from untreated and TNFα-treated and oxLDL-treated groups. Each experimental point is an average of the technical replicates. B) Volcano plot of DEPs in EVs from TNFα-treated HUVEC compared to proteins in EVs from untreated cells. C) Volcano plot of DEPs in EVs from oxLDL-treated HUVEC compared to proteins in EVs from untreated cells. D) Volcano plot of DEPs in EVs from TNFα-treated HUVEC compared to proteins in EVs from oxLDL-treated cells. Significantly different proteins were determined as those with an adjusted *p*-value <0.05 and a fold change >2 (red dots). Data are from three biological replicates each measured in triplicate. E) Heat map of the top 10 significant DEPs in EVs from TNFα-treated cells compared to EVs from untreated cells ranked by log2FC. F) Heatmap for the top 10 significant DEPs in EVs from TNFα-treated cells compared to EVs from oxLDL-treated cells ranked by log2FC.

A total of 386 proteins were significantly up- or down-regulated in EVs derived from TNFα-treated HUVEC compared to EVs from untreated cells based on a false discovery rate (FDR) < 0.05 and fold change > 2 (Figure 3B and Supplemental Data File 3). Upregulated proteins included E-selectin (SELE) (log2FC = 4.79; *p* = 1.92 × 10^-7^), PRAMEF2 (log2FC = 8.46, *p* = 3.5 × 10^-3^), and dual specificity phosphatase 7 (DUSP7) (log2FC = 4.38, *p* = 8.5 × 10^-4^). Only one protein, H1.4 linker histone cluster member (HIST1H1E) (log2FC = 2.42, *p* = 0.032), was found to be significantly differentially expressed in EVs from HUVECs treated with oxLDL compared to EVs from untreated HUVECs (Figure 3C and Supplemental Data File 3).

The comparison of proteomes between EVs derived from TNFα- and oxLDL-treated HUVECs was performed to investigate how distinct pathological stimuli that represent different mechanisms of endothelial activation or injury effect the proteome of ECdEVs. TNFα is a potent pro-inflammatory cytokine known to induce endothelial apoptosis, while oxLDL is associated with oxidative stress and early endothelial activation without triggering apoptosis at the concentration used in the study. This comparison allowed us to identify proteins selectively packaged into EVs during endothelial apoptosis (TNFα) versus non-apoptotic activation (oxLDL). EVs derived from TNFα-treated cells significantly and differentially expressed 147 proteins compared to EVs from oxLDL-treated HUVEC, with XIRP1 showing the greatest difference in log2FC (log2FC = 4.74, *p* = 3.19 × 10^-2^), followed closely by E-selectin (log2FC = 4.51, p = 4.11 × 10^-7^) and DUSP7 (log2FC = 4.35, p = 3.83 × 10^-3^) (Figure 3D, Supplemental Data File 3). Heatmaps of the top ten candidate biomarkers in TNFα-treated vs. untreated EVs (Figure 3E) and TNFα-treated vs. oxLDL EVs (Figure 3F) show DEPs between treatments based on the greatest log2FC. Specifically, in Figure 3E the proteins are ranked in order related to highest to lowest expression of these proteins in EVs from TNFα-treated vs. untreated cells, whereas in Figure 3F the proteins are ranked in order related to highest to lowest expression of these proteins in EVs from TNFα-treated vs. oxLDL-treated cells. E-selectin expression is increased in both Figure 3E and 3F because E-selectin was only increased in EVs from TNFα-treated HUVEC and therefore shows as increased expression in both TNFα vs. untreated and TNFα vs. oxLDL groups. The biological functions of the top ten candidate biomarkers in TNFα-treated vs. untreated EVs are shown in Table 1 [50–61]. Log2FC and adjusted *p*-values for the top ten candidate biomarkers for TNFα vs. untreated EVs (Table 2A) and TNFα vs. oxLDL EVs (Table 2B) are provided. Interestingly, three proteins (E-selectin, DUSP7 and PRAMEF2) were in the top ten DEPs in EVs from both the TNFα-treated vs. untreated cells and TNFα-treated vs. oxLDL-treated cells (Tables 2A&B).

**Table 1.** Functions of the top ten significantly and differentially expressed proteins in EVs derived from TNFα-treated vs. untreated HUVEC.

| Gene Abbreviation | Gene name | Function | References |
| --- | --- | --- | --- |
| PRAMEF2 | PRAME family member 2 | Part of the cullin 2 E3 ubiquitin ligase complex | Ghosh & Das 2021 [50] |
| TXNL1 | Thioredoxin like 1 | Reduces disulphide bonds in proteins. Acts as an ATP-independent chaperone protein | Andor *et al.* 2023 [51] |
| AP2A1 | Adaptor related protein complex 2 subunit alpha 1 | An adaptor protein that links membranes to clathrin lattices during endocytosis | Jackson *et al.* 2010 [52] |
| SELE | E-selectin | Endothelial adhesion molecule expressed in response to inflammatory cytokines. Mediates leukocyte adhesion to the vascular endothelium by interacting with P-selectin glycoprotein ligand 1 on leukocytes | Rahman *et al.* 1998 [53]; Laviola *et al.* 2013 [54] |
| MMP3 | Matrix metalloproteinase 3/stromelysin 1 | Degrades the extracellular matrix during tissue remodelling. Treatment of HUVEC with TNFα increased MMP3 mRNA and protein expression | Hanemaaijer *et al.* 1993 [55] |
| MYLIP | Myosin regulatory light chain interacting protein | E3 ubiquitin protein ligase that ubiquitinates myosin regulatory light chain and LDL receptor leading to their degradation | Bornhauser *et al.* 2003 [56]<br>Zelcer *et al.* 2009 [57] |
| DUSP7 | Dual specificity phosphatase 7 | Protein serine/threonine/tyrosine phosphatase that dephosphorylates ERK1/2 | Caunt & Keyse 2013 [58] |
| SDPR | Serum deprivation protein response | Regulates curvature of membranes and caveolae structure | Hansen *et al.* 2009 [59] |
| LRRC59 | Leucine rich repeat containing 59 | A membrane-associated protein that is localised to the endoplasmic reticulum. Transports fibroblast growth factor 1 into the nucleus | Zhen *et al.* 2012 [60] |
| GABPB2 | GA binding protein transcription factor subunit beta 2 | ETS transcription factor that regulates haematopoietic stem cell survival | Yu *et al.* 2012 [61] |

**Table 2.**
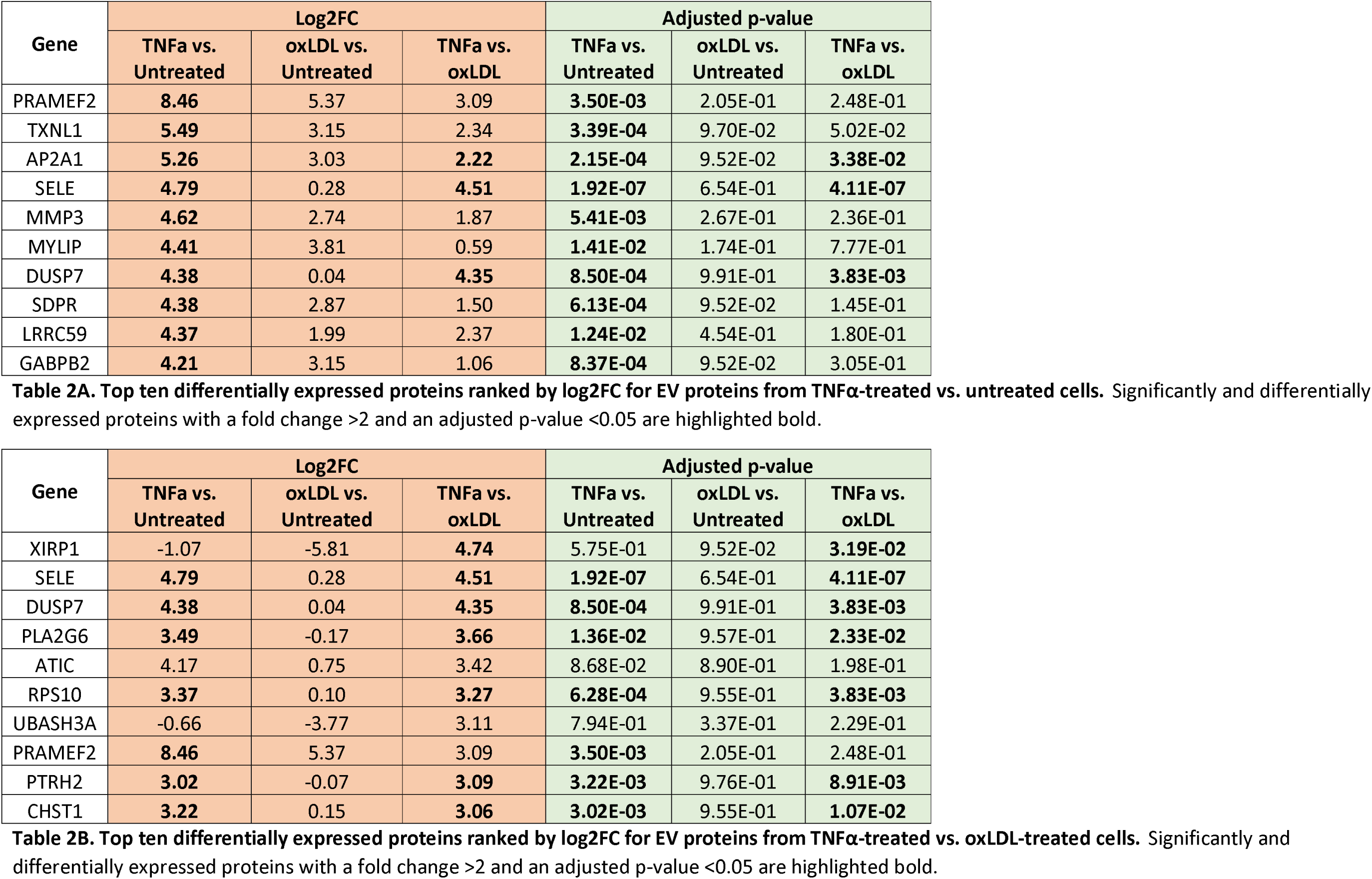
Top ten differentially expressed proteins ranked by log2FC for EV proteins from A) TNFα-treated vs. untreated cells and B) TNFα-treated vs. oxLDL-treated cells. Significant DEPs with a fold change >2 and an adjusted *p*-value <0.05 are highlighted bold.

### Protein-protein network analysis and GO-term enrichment analysis

Protein-protein association network analysis of significant DEPs present in EVs isolated from TNFα-treated cells compared to EVs from untreated HUVEC was performed using the STRING protein-protein interaction database [49]. This showed more protein-protein interactions than expected from a random set of proteins, with 215 edges compared to the expected 125, and a protein-protein interaction enrichment *p*-value of 1.62 × 10^-13^. This analysis revealed that, compared to EVs from untreated cells, EVs from TNFα-treated HUVEC were enriched in proteins involved in translation (RPL proteins, EIF5A), proteosome-mediated ubiquitin-dependent protein catabolism (PSMA6, PSMD2, PSMC3 etc.), Rho/Ras signal transduction (RHOA, RALA, RALB, AKAP13, GNA12, ARHGDIB), translational regulation (YBX1, TNRC6A, DHX9, SYNCRIP), and leukocyte cell-cell adhesion (SELE, VCAM1, CD40, ICAM1, ICAM2) (Figure 4A). GO-term enrichment analysis revealed biological process terms such as regulation of plasma membrane repair, mRNA stabilisation, negative regulation of tyrosine kinase signalling, positive regulation of translation, and cell spreading (Figure 4B). Cellular component term analysis revealed that proteins associated with sorting endosomes, flotillin complexes, tetraspanin-enriched microdomains and proteosome complexes were enriched in EVs from TNFα-treated HUVEC compared to EVs from untreated cells (Figure 4C).

**Figure 4.**
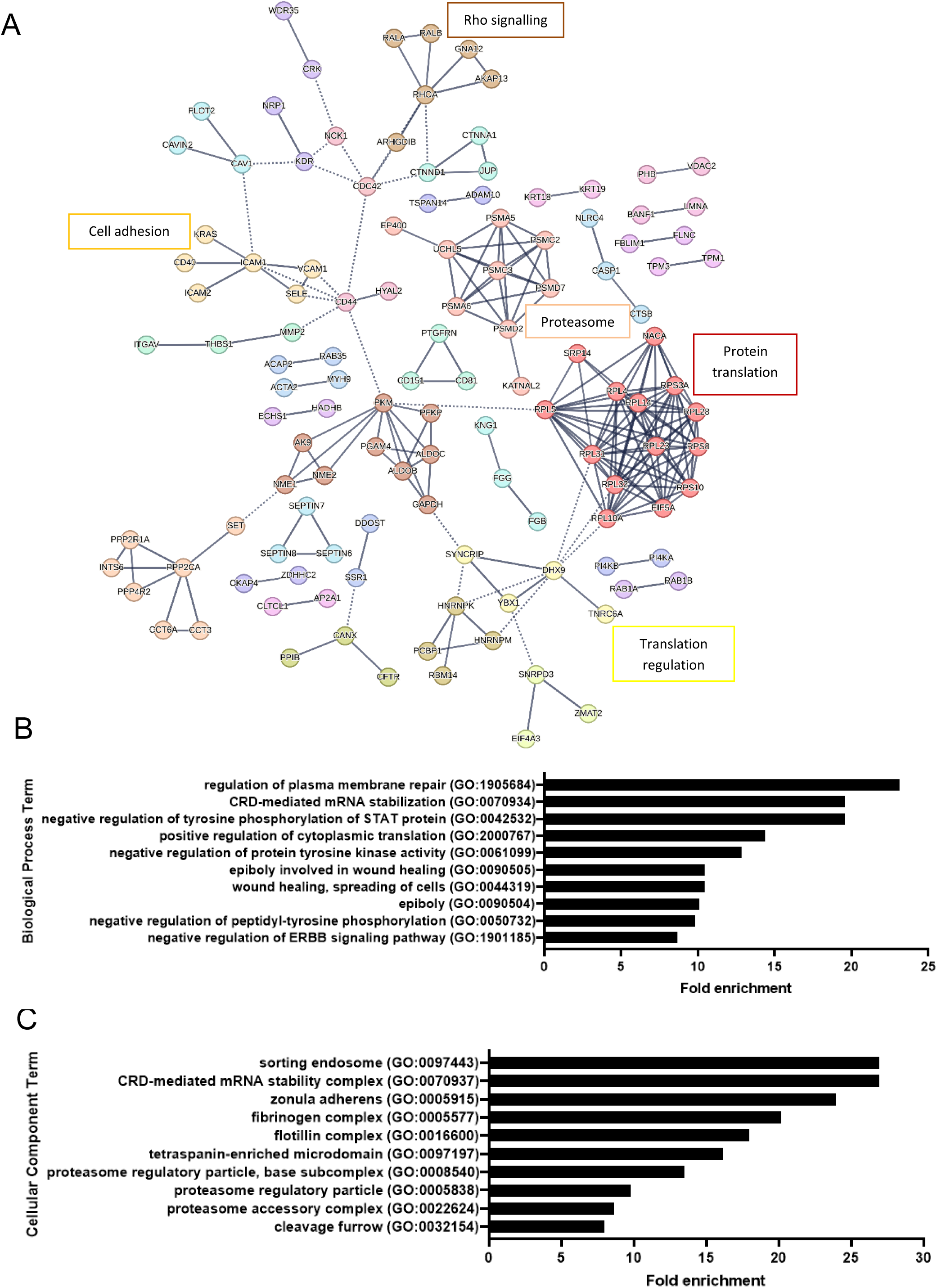
STRING and GO analysis of proteins significantly and differentially expressed in HUVEC-derived EVs from untreated vs. TNFα-treated cells. Proteins that were significantly and differentially expressed in EVs from untreated and TNFα-treated HUVEC compared to EVs from untreated cells were analysed using A) STRING analysis to examine protein-protein interaction networks showing network edges as confidence (line thickness indicates strength of data to support interactions). PANTHER gene ontology overrepresentation test with an FDR of p<0.05 was used to examine enrichment of B) biological process terms and C) cellular component terms associated with significant DEPs in EVs derived from TNFα-treated cells compared to EVs from untreated cells.

Proteins significantly and differentially expressed in EVs from HUVEC treated with TNFα compared to EVs from oxLDL-treated cells showed 33 edges compared to the expected 18, and a protein-protein interaction enrichment *p*-value of 0.0009 (Figure 5A). These included proteins involved in protein translation (RPL proteins), Rho signalling (RHOA, ARHGDIA, ARHGDIB), nucleotide metabolism (AK9, NME1, NME2), and regulation of translation (YBX1, SYNCRIP, DHX9). GO-term enrichment analysis revealed biological processes terms such as protein localisation to cytoplasmic stress granule, vascular endothelial growth factor signalling, regulation of cell-substrate adhesion, negative regulation of cell motility, and regulation of plasma membrane bounded cell projection organisation (Figure 5B). Cellular component term analysis revealed proteins associated with mRNA stability complexes, sorting endosomes, cortical cytoskeleton and adherens junctions (Figure 5C).

**Figure 5.**
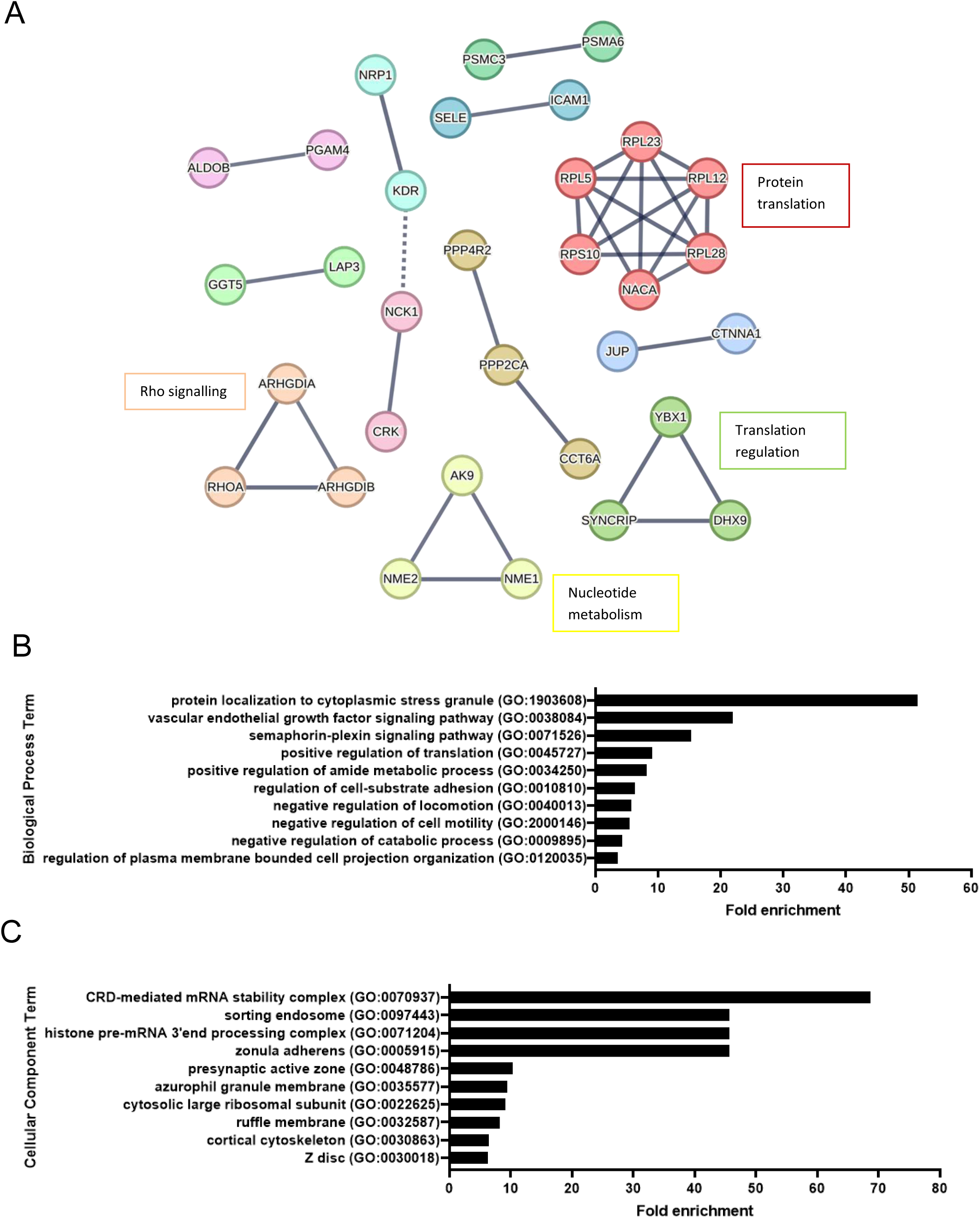
STRING and GO analysis of proteins significantly and differentially expressed between HUVEC-derived EVs from TNFα-treated vs. oxLDL-treated cells. Proteins that were significantly and differentially expressed in EVs from TNFα-treated and oxLDL-treated HUVEC were analysed using A) STRING analysis to examine protein-protein interaction networks showing network edges as confidence (line thickness indicates strength of data to support interactions). PANTHER gene ontology overrepresentation test with an FDR of p<0.05 was used to examine enrichment of B) biological process terms and C) cellular component terms associated with significant DEPs in EVs derived from TNFα-treated compared to EVs from oxLDL-treated cells.

## Discussion

Endothelial cells release EVs in response to vascular injury, inflammation and endothelial dysfunction and have been suggested as potential biomarkers of CVD. Identifying key proteins that are packaged into EVs by endothelial cells in response to various stimuli relevant to CVD may provide biomarkers that reveal information about the state of the vascular endothelium in disease conditions. The aim of this study was to examine the proteome of endothelial EVs released *in vitro* in response to stimuli that endothelial cells may be exposed to in patients with CVD (Figure 6). Endothelial cells were treated with stimuli that are known to increased EV release and assessed whether this was associated with apoptosis.

**Figure 6.**
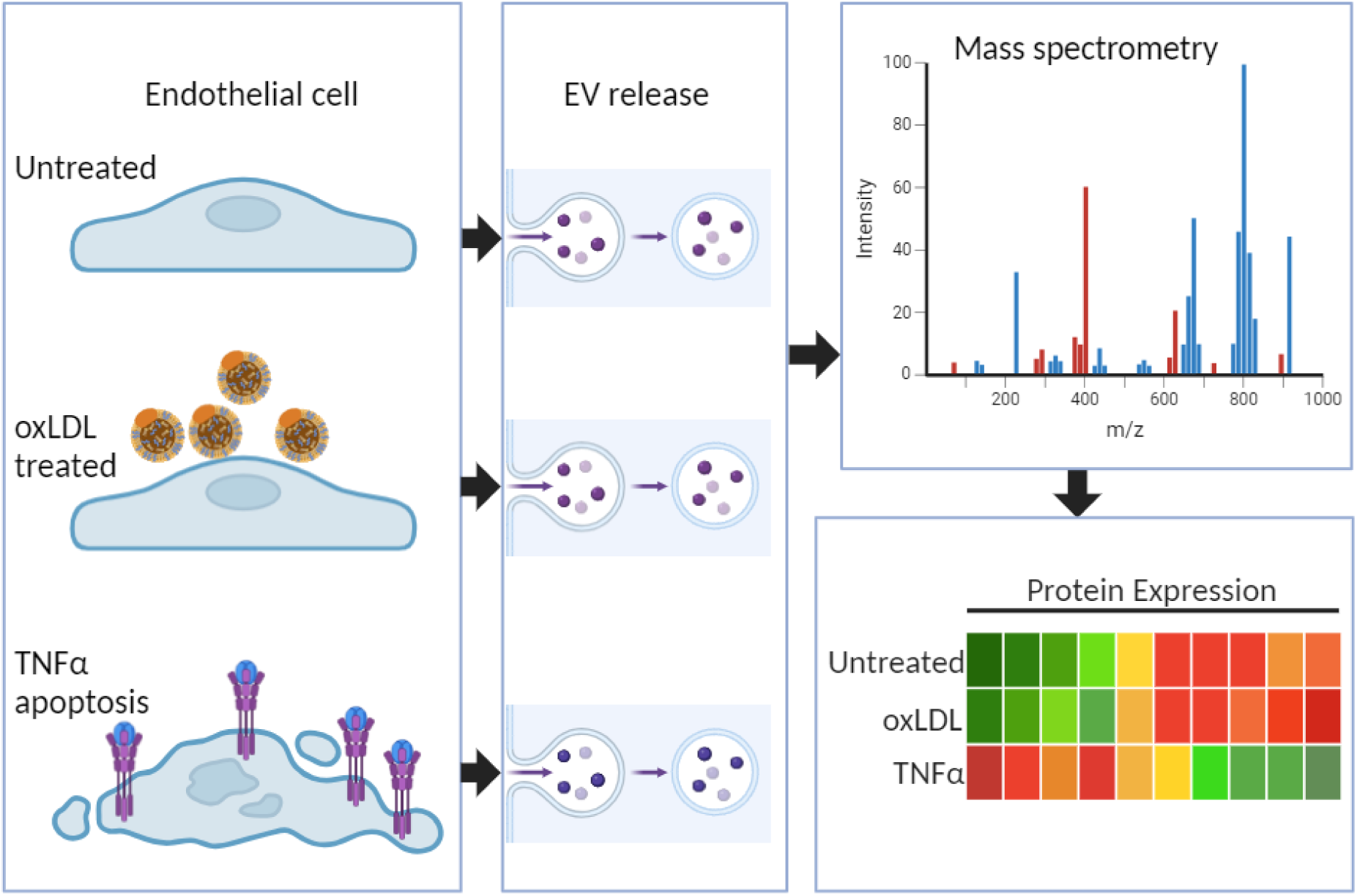
A schematic diagram showing treatment of endothelial cells with TNFα or oxLDL results in the release of EVs with different protein cargos. Created in BioRender. Collier, M. (2025) https://BioRender.com/d18z861

In this study endothelial cells were treated with TNFα because levels of this proinflammatory cytokine have been shown to be elevated in the blood of patients with CVD [31,32,33] and TNFα directly affects the vascular endothelium, resulting in vascular inflammation and reduced vasodilation, which are characteristics of endothelial dysfunction [34,35,36]. Specifically, TNFα downregulates the expression of eNOS [62] leading to reduced nitric oxide synthesis and reduced vasodilation.

TNFα also induces the expression of adhesion molecules on the surface of endothelial cells such as E-selectin, VCAM and ICAM which promote monocyte adhesion and transmigration [53,54,63]. TNFα is commonly used at a concentration of 10 ng/ml to stimulate endothelial cells *in vitro*. For example, Kuldo *et al*. (2005) [43] showed the upregulation of the expression of various adhesion molecules and cytokines in HUVEC stimulated with 10 ng/ml TNFα for 6 h. Hosseinkhani *et al*. (2020) [44] treated HUVEC with 10 ng/ml TNFα for 24 h to induce EV release, whereas Voisard *et al*. (1998) [45] demonstrated that 10 ng/ml TNFα significantly reduced HUVEC proliferation compared to untreated cells. In agreement with previous studies, we observed that TNFα induced apoptosis in HUVEC [64] (Figure 1) and increased the release of EVs from these cells [8] (Figure 2).

We also examined the influence of oxLDL on EV release from endothelial cells. Increased oxLDL levels have been detected in patients with CVD [37,38]. oxLDL directly affects endothelial cells by inducing endothelial injury and apoptosis through increased oxidative stress [39]. Incubation of endothelial cells with oxLDL also activates CD40/CD40L signalling in endothelial cells [41], increases the expression of adhesion molecules [40,41], and inhibits eNOS activity [42], promoting leukocyte adhesion and reducing vasodilation. The effects of TNFα and oxLDL on the proteome of ECdEVs were therefore assessed in this study because of their importance in the progression of CVD through their role in promoting various aspects of endothelial dysfunction. Previous studies have shown that high concentrations of oxLDL (≥25 µg/mL) induce apoptosis in HUVEC [39,65]. Here, using a lower concentration of oxLDL (10 µg/mL), we did not observe endothelial apoptosis (Figure 1), which is in line with previous findings in coronary artery endothelial cells (HCAECs) [66]. However, lower concentrations of oxLDL (≤20 µg/mL) have been shown to activate endothelial cells resulting in increased cell surface expression of P-selectin [40] and CD40 [41], and ROS production [46]. We observed a non-significant increase in CD40 expression on EVs released from oxLDL-treated cells vs. EVs from untreated cells (log2FC = 1.15, p = 4.68 × 10^-1^). This, in addition to the non-significant increase in the levels of EVs released in response to a lower concentration of oxLDL than previously reported [13], may therefore represent the effects of lower levels of oxLDL in earlier stages of atherosclerosis. In contrast, EVs from TNFα-treated cells may represent a later phase, or more established disease, which is accompanied by apoptosis.

EV released from the HUVECs were isolated from the conditioned media using high speed centrifugation at 40,000 *g*. This likely resulted in the isolation of both large and small (exosome-like) EVs. This was supported by the fact that the mass spectrometry analysis revealed the presence of many EV-associated proteins listed in the ExoCarta and Vesiclepedia databases, such as tetraspanins CD9 and CD81, heat shock proteins, Alix, annexins and flotillins. Although there are no EV markers that definitely distinguish between exosomes and microvesicles, proteomics analysis of the isolated EVs revealed the presence of the tetraspanins CD81 and CD9 and the protein Alix which is involved in exosome biogenesis. In fact, EVs from TNFα-treated HUVEC were enriched for the EV marker CD81 compared to EVs from untreated cells, further confirming the isolation of EVs and highlighting potential differences in EV populations between treatments. Proteomics analysis of the isolated EVs also revealed the presence of annexin A1, which has been shown to be present in microvesicles shed from the cell surface but not present in exosomes [67], further indicating a mixed population of EVs was isolated by high-speed centrifugation. The isolation of EVs was further confirmed by GO-term enrichment analysis of proteins detected in EVs from TNFα-treated cells which showed enrichment of cellular component terms related to EV formation including endosome sorting, flotillin complex and tetraspanin-enriched microdomain in EVs (Figure 4C).

Although several previous studies have examined of the proteome of EVs from TNFα-treated endothelial cells compared to untreated cells [27,29,30,68,69] we detected a total of 1355 quantifiable proteins in HUVEC-derived EVs, representing a much-improved sensitivity. In agreement with previous studies, EVs from TNFα-treated cells showed extensive differences in their protein composition compared to EVs from untreated cells [68,69], but also in comparison to EVs from oxLDL-treated cells, which has not been examined previously. The protein content of EVs released from oxLDL-treated HUVEC was also distinct but was more similar to the proteome of EVs from untreated cells than EVs from TNFα-treated cells, as shown by both the PCA (Figure 3A) and the heatmap of DEPs (Figure 3E).

In particular, the endothelial adhesion molecule E-selectin (SELE; CD62e) was significantly increased in EVs derived from HUVEC treated with TNFα compared to EVs from untreated or oxLDL-treated cells. E-selectin is an endothelial cell adhesion molecule which mediates endothelial cell-leukocyte interactions and is upregulated by TNFα [53,54]. Increased expression of E-selectin is a well-known marker of endothelial cell activation, dysfunction and apoptosis, and increased E-selectin expression has been detected on endothelial cells in CVD [70,71]. E-selectin has previously been detected by flow cytometry on EVs released from TNFα-stimulated endothelial cells *in vitro* [18,72], and clinical studies have detected increased levels of E-selectin-positive EVs in the circulation of patients with CVD [25,73,74] and diseases associated with increased risk of CVD such as chronic kidney disease [75] and chronic obstructive pulmonary disease [76]. Amabile *et al.* (2009) [25] detected increased levels of circulating E-selectin-positive endothelial-derived EVs in patients with pulmonary hypertension in patients with a poor outcome and suggested that levels of these EVs could be used as a biomarker of CVD and to stratify patients for treatment.

Surprisingly, previous proteomics studies of EVs generated *in vitro* from TNFα-treated endothelial cells have not detected increased levels of E-selectin [27,29,68,69]. E-selectin is rapidly expressed in HUVEC in response to TNFα, with protein expression peaking at 6 h post-treatment and returning nearer to baseline levels by 24 h [63]. The lack of E-selectin in ECdEVs in previous *in vitro* studies may therefore be due to differences in the time point of EV isolation between studies, since we used a relatively short treatment of 6 h, rather than 24 h stimulation used in many past studies. This may also explain why we did not observe significant increases in markers of cellular activation in EVs from cells incubated with oxLDL. For example, 24-48 h incubation with 12.5 µg/ml oxLDL was required for significant CD40 expression on the surface of HUVEC [77]. Therefore, this is the first *in vitro* study to show increased levels of E-selectin on EVs from TNFα-treated HUVEC using mass spectrometry. This is important because E-selectin is a preferred biomarker for ECdEVs in the circulation, and our data shows an association between E-selectin-positive ECdEVs and endothelial cell apoptosis, indicating that E-selectin could be a marker of endothelial damage and apoptosis in CVD, although further mechanistic studies would be required to confirm this.

Another protein that was significantly increased in EVs in response to TNFα-treatment was dual specificity phosphatase 7 (DUSP7). This enzyme is localised to the cytoplasm where it dephosphorylates ERK1/2 of the mitogen activated protein kinase (MAPK) pathway [58]. A previous study has shown that treatment of HUVEC with inflammatory cytokines including TNFα, results in the upregulation of DUSP7 [78]. Interestingly, DUSP7 supresses the secretion of TNFα from endothelial cells by dephosphorylating and therefore inactivating ERK1/2 leading to reduced TNFα expression [79]. This was further confirmed in a study by Leng *et al.* (2018) [80] which showed that homocysteine-induced degradation of DUSP7 resulted in increased ERK1/2 phosphorylation and upregulation of TNFα expression in HUVEC. The upregulation of DUSP7 in HUVEC-derived EVs in response to TNFα therefore suggests a negative feedback mechanism in endothelial cells to reduce levels of TNFα.

GO-term enrichment analysis of DEPs in EVs derived from TNFα-treated HUVEC compared to DEPs in EVs from untreated cells revealed enrichment of biological process terms such as regulation of plasma membrane repair, coding region instability determinant (CRD)-mediated mRNA stabilisation, positive regulation of cytoplasmic translation, and negative regulation of tyrosine kinase activity.

Enrichment of DEPs involved in plasma membrane repair may be related to the processes of EV generation and release, since membrane repair involves similar cellular mechanisms as those required for EV release such as exocytosis and rearrangement of the actin cytoskeleton [81]. EVs from TNFα-treated HUVEC also showed enrichment of DEPs associated with the cellular component terms sorting endosome, flotillin complex, and tetraspanin-enriched microdomain, which are associated with EV biogenesis. Both GO-term enrichment and STRING analysis revealed enrichment of biological process terms and protein clusters involved in protein translation, which may reflect the activation of intracellular signalling pathways within the endothelial cells in response to TNFα, leading to altered mRNA stability and changes in protein expression. Interestingly, several enriched biological process terms were associated with negative regulation of protein tyrosine kinase activity and related pathways such as STAT phosphorylation and ErbB signalling.

Tyrosine kinases are activated by TNFα-signalling in endothelial cells and lead to increased expression of adhesion molecules [82] and apoptosis [83]. Enrichment of DEPs associated with negative regulation of tyrosine kinase activity may therefore indicate activation of signalling pathways in the endothelial cells to regulate TNFα-mediated apoptosis.

GO-term enrichment analysis of DEPs in EVs derived from TNFα-treated HUVEC compared to DEPs in EVs from oxLDL-treated cells revealed enrichment of biological processes associated with response to cellular stress and changes in cell signalling pathways, such as protein localisation to stress granules [84] and VEGF signalling [85]. GO-term enrichment analysis also revealed enrichment of biological process terms involved in the negative regulation of cell migration. This was further confirmed by STRING analysis of DEPs which showed enrichment of Rho associated-proteins that mediate cell movement, suggesting changes in cell motility in cells treated with TNFα. This further demonstrates how the proteomic profile of EVs provides information on the cellular processes occurring in the parent cells. Furthermore, EV-mediated transfer of these proteins released from apoptotic or injured endothelial cells could affect surrounding cells and propagate endothelial dysfunction in patients with CVD.

A limitation of this study is that HUVEC are foetal endothelial cells derived from veins and are not necessarily representative of endothelial cells in adult arteries which may differ in the release of EVs and protein loading into EVs in response to the stimuli used in this study. Furthermore, the serum-free conditions used to prevent interference from bovine-derived EVs present in FCS resulted in increased background levels of apoptosis and also restricted the study to a 6 h incubation prior to EV isolation, which may have limited the detection of proteins that require longer time periods for upregulation of gene expression. Another consideration is that high-speed centrifugation at 40,000 *g* was used to isolate EVs from the conditioned media. This centrifugation speed has been used in previous studies to isolate EVs from urine [86] and cell culture media [87]. Jeppesen *et al.* (2014) [88] demonstrated centrifugation of cell culture media at a lower centrifugation speed of 33,000 *g* resulted in the isolation of both small (up to 100 nm) and large (100-300 nm) EVs.

Furthermore, Durak-Kozica *et al*. (2022) [89] reported that isolation of endothelial cell-derived EVs from culture media resulted in a mixed population of both microvesicles and exosomes even at a lower centrifugation speed of 18,000 *g*. It should also be noted that the characterisation of EVs in the HUVEC-conditioned media using NTA may not be representative of the size distributions of EVs isolated for proteomics analysis by high-speed centrifugation. However, proteomics analysis of the isolated EVs revealed enrichment of EV-associated proteins, indicating that EVs were efficiently isolated using this protocol. Finally, E-selectin-positive EVs have previously been suggested as a marker of endothelial dysfunction or injury in patients with CVD [25,73]. It would therefore be interesting to validate if any of the other top ten candidate biomarkers detected in this study are also differentially expressed in EVs from patients with CVD compared to EVs from healthy controls.

In this study LC-MS/MS was used to examine the proteome of EVs due to its high sensitivity, high resolution, and dynamic range, particularly for complex samples such as EVs. It enables both the identification and quantification of a wide range of proteins with a relatively low sample input, which is critical in EV research. However, we acknowledge that other mass spectrometry techniques such as secondary ion mass spectrometry (SIMS) have been successfully used to examine the proteome of EVs [90]. SIMS provides valuable spatial and surface composition data, although its application to EV proteomics is still limited compared to LC-MS/MS. Another recent study by Kasprzyk-Pochopień *et al.* (2025) [91] compared nanoLC-MALDI-MS/MS and nanoLC-TIMS-MS/MS for the proteomic analysis of EVs and concluded that nanoLC-TIMS-MS/MS detected significantly more proteins than nanoLC-MALDI-MS/MS, highlighting how increased sensitivity of mass spectrometry techniques can improve EV protein detection.

In conclusion, this study has further confirmed that the protein content of EVs released from endothelial cells is significantly altered in response to apoptotic stimuli such as TNFα [18,69]. EV protein profiles also differed between treatment of HUVEC with TNFα and a low-level of oxLDL, suggesting that different stimuli relevant to CVD result in selective packaging of proteins into EVs (Figure 6). Ten potential biomarkers were identified in this study which distinguished between EVs released from TNFα-treated, apoptotic endothelial cells and untreated cells or cells treated with oxLDL. Due to significant increases in E-selectin-positive EVs released from TNFα-treated cells, and the high abundance of this protein compared to levels in EVs from other treatment groups, this suggests that E-selectin could be used as a potential candidate biomarker for endothelial damage and apoptosis in CVD.

## Author contributions

MEWC designed the study, conducted the cell experiments and wrote the manuscript. AHG oversaw the design and execution of the study and co-wrote the manuscript. JKS and PAQ conducted the LC-MS/MS analysis and THC analysed the mass spectrometry data. DJLJ oversaw the LC-MS/MS analysis. All authors read and approved the manuscript.

## Data Availability Statement

All data is available on reasonable request.

