## Supplemental Data File 1 for "Proteomic analysis of extracellular vesicles released from endothelial cells *in vitro* reveals increased levels of E-selectin and dual specificity phosphatase 7 as a potential marker of TNFα-mediated apoptosis"

**Supplemental Data File 1. The gradient in 110 minutes of solvent A and B used in LC ESI-MS/MS analysis**

ACE Experimental Record

Inlet Method File: c:\projects\pq_new_may_2014.pro\acqudb\110min_withacnramp

---------------------	Run method parameters	----------------

-- PUMP --

Waters GI Pump

1 --------------------------------------------

Application Mode: Single Pump Trapping

Pump Type: BSM1

Run Time: 110.00 min

Solvent Selection A: A1

Solvent Selection B: B1

Seal Wash: 30.0 min

Switch 1: No Change

Switch 2: No Change

Switch 3: No Change

Chart Out 1: System Pressure

Chart Out 2: %B

Run Events: Yes

[Gradient Table]

Time(min) Flow Rate(uL/min) %A %B Curve

1. Initial 0.300 97.0 3.0

2. 20.00 0.300 86.0 14.0 6

3. 30.00 0.300 80.0 20.0 6

4. 40.00 0.300 75.0 25.0 6

5. 51.00 0.300 69.0 31.0 6

6. 52.00 0.300 69.0 31.0 6

7. 52.10 0.300 69.0 31.0 6

8. 52.20 0.300 69.0 31.0 6

9. 53.00 0.300 65.0 35.0 6

10. 53.10 0.300 65.0 35.0 6

11. 54.00 0.300 63.0 37.0 6

12. 55.00 0.300 58.0 42.0 6

13. 63.00 0.300 31.0 69.0 6

14. 65.00 0.300 97.0 3.0 6

15. 80.00 0.300 50.0 50.0 6

16. 80.50 0.300 10.0 90.0 6

17. 82.20 0.300 97.0 3.0 6

18. 87.50 0.300 97.0 3.0 6

19. 99.50 0.300 50.0 50.0 6

20. 101.50 0.300 10.0 90.0 6

21. 103.50 0.300 97.0 3.0 6

22. 110.00 0.300 97.0 3.0 6

Analytical Low Pressure Limit: 0 psi

Analytical High Pressure Limit: 10000 psi

Sample Loading Time: 3.00 min

Trapping Flow Rate: 5.000 uL/min

Trapping %A: 99.5

Trapping %B: 0.5

Trapping Low Pressure Limit: 0 psi

Trapping High Pressure Limit: 10000 psi

Flow Rate A Data Channel: No

Flow Rate B Data Channel: No

Solvent Name A: Water

Solvent Name B: Acetonitrile

Comment:

System Pressure Data Channel: Yes

Flow Rate Data Channel: No

%A Data Channel: No

%B Data Channel: No

Primary A Pressure Data Channel: No

Accumulator A Pressure Data Channel: No

Primary B Pressure Data Channel: No

Accumulator B Pressure Data Channel: No

Degasser Pressure Data Channel: No

3 --------------------------------------------

Run Time: 110.00 min

Pump A...

Aux Pump Role: Auxiliary

Aux Solvent Name: Water

Aux Flow Rate: 0.500 uL/min

Aux Solvent Selection: A1

Aux Low Pressure Limit: 0 psi

Aux High Pressure Limit: 10000 psi

Pump B...

Aux Pump Role: Lock Mass

Aux Solvent Name: Water

Aux Flow Rate: 0.500 uL/min

Aux Solvent Selection: B1

Aux Low Pressure Limit: 0 psi

Aux High Pressure Limit: 10000 psi

-- END PUMP --

-- AUTOSAMPLER --

Waters Acquity AutoSampler

Run Time: 110.00 min

Comment:

Loop Option: Partial Loop

LoopOffline: Disable

Weak Wash Solvent Name: Water

Weak Wash Volume: 600 uL

Strong Wash Solvent Name: Acetonitrile

Strong Wash Volume: 200 uL

Target Column Temperature: 40.0 C

Column Temperature Alarm Band: Disabled

Target Sample Temperature: 8.0 C

Sample Temperature Alarm Band: Disabled

Full Loop Overfill Factor: Automatic

Syringe Draw Rate: Automatic

Needle Placement: 0.5

Pre-Aspirate Air Gap: Automatic

Post-Aspirate Air Gap: Automatic

Column Temperature Data Channel: No

Ambient Temperature Data Channel: No

Sample Temperature Data Channel: No

Sample Pressure Data Channel: No

Switch 1: No Change

Switch 2: No Change

Switch 3: No Change

Switch 4: No Change

Chart Out: Sample Pressure

Sample Temp Alarm: Disabled

Column Temp Alarm: Disabled

Run Events: Yes

SampleLoop: 5.000

Saved as Trizaic: No

nanoTile Cool Down: 2.1

Sample Run Injection Parameter

Injection Volume (ul)	-	1.00

-- END AUTOSAMPLER --

---------------------------- oOo -----------------------------

End of experimental record.

------------------- Waters Acquity SM Postrun Report ---------------

Software Version: 1.42.1248

Firmware Version: 1.42.292 (Mar 09 2011)

Checksum: 0x5bf322dd

Serial Number: M09NPS661M

Sample Syringe Size: 100.0

Sample Loop Size: 5.0

NeedleType: PEEK

Minimum Sample Temperature: 0.0

Maximum Sample Temperature: 0.0

Average Sample Temperature: 0.0

Minimum Column Temperature: 36.5

Maximum Column Temperature: 40.1

Average Column Temperature: 0.0

Measured Loop Volume: 5.500

Measured Loop Volume No Pressure: 5.560

---------------------------- oOo -----------------------------

------------------- Waters GI Pump Postrun Report ---------------

---------------------------- oOo -----------------------------
